# The effect of meal temperature on heart rate in *Rhodnius prolixus*

**DOI:** 10.1101/685305

**Authors:** Chloé Lahondère, Maurane Buradino, Claudio R. Lazzari

**Author notes:** To whom all correspondence should be addressed: Chloé Lahondère, Department of Biochemistry, Virginia Polytechnic Institute and State University, Blacksburg, VA, 24061, USA.

## Abstract

*Rhodnius prolixus* is able to cool down the ingested blood during feeding on a warm-blooded host. This is possible because of a counter-current heat exchanger located in its head, which transfers heat from the warm blood to the insect haemolymph and can dissipate through the head cuticle. Given the key role haemolymph circulation in thermoregulation, we investigated the modulation of the activity of the heart during the warmed meal intake. We evaluated the impact of meal temperature on the heart rate and found that feeding led to an increase in the frequency of heart contractions, which increases with increasing food temperature. We also found that females have a higher heart rate during feeding compare to males.

**HIGHLIGHTS:**

- Feeding increases the heart rate of *Rhodnius prolixus*
- The higher the meal temperature, the higher the heart rate becomes
- Females have a higher heart rate than males

## INTRODUCTION

Despite its simplicity, the open circulatory system of insects plays a paramount role in the transport of hemolymph and other molecules. The mechanical impulse maintaining the haemolymph in movement is mainly provided by the heart. Its structure in *Rhodnius prolixus* is particular, in the sense that haemolymph is only gathered at the very end of the abdominal cavity, where four pairs of ostia and alary muscles are located, to impulse it into the head (Chiang *et al.*, 1990). Hemolymph carries nutrients, immune cells, waste products and substances such as hormones (Jones, 1964). Among these molecules, the biogenic amine serotonin (5-hydroxytryptamine) is implicated in numerous physiological processes and its action has been extensively studied in *R. prolixus*. Serotonin acts both as a neuromodulator, delivered by nerve supply and as a neurohormone transported *via* the hemolymph. Serotonin secretion is timely related to feeding-associated events (Orchard, 2006) such as diuresis and excretion (*i.e.*, diuretic hormone) (Maddrell *et al*., 1969; Maddrell *et al*., 1991; Martini *et al*., 2007), plasticization of the cuticle (Ianowski *et al*., 1998; Orchard *et al*., 1988; Reynolds, 1974), muscle contractions of the salivary glands (Orchard and Te Brugge, 2002), crop (Farmer *et al*., 1981) and the hindgut (Orchard *et al*., 1988). In addition, serotonin increases the contractions of the dorsal vessel of *R. prolixus* both *in vivo* and *in vitro* (Chiang *et al*., 1992) and its concentration in hemolymph also increases during feeding (Lange *et al*., 1989). More recently, it has been shown that CCAP (Crustacean CardioActive Peptide) is also involved in the control of the heart rate in this species (Lee *et al.*, 2013). We can thus expect, the dorsal vessel contractions to increase during feeding.

The circulatory system of *R. prolixus* is involved in thermoregulation, as it is in some other insects including bees, bumblebees and moths (Heinrich, 1976, 1987, 1993; May, 1979). *R. prolixus* bugs have a sophisticated thermoregulatory mechanism (*i.e.* counter current heat exchanger located in the head), which helps them maintaining a regional heterothermy during feeding on warm-blooded hosts and minimizes overheating due to the rapid ingestion of warm blood (Lahondère *et al.*, 2017). By reducing the risk of thermal stress, this mechanism protects their physiological integrity (Benoit *et al.*, 2019). Moreover, cooling down during blood-feeding, allows these bugs to reduce the risk of being victim of cannibalism (Lazzari *et al.*, 2018). Indeed, as heat is an important host-seeking cue that elicits biting, indiviuals with an abdomen full of warm blood could be a target for conspecifics to obtain an “easy” blood-meal without taking the risk of being exposed to the host’s anti-parasitic behaviors.

Given the role of the circulatory system in thermoregulation during feeding, it is worth exploring to what extent the temperature of the ingested blood regulates the activity of the heart, in order to maintain an adequate heat transfer along the insect body during feeding. In this context, the present study examined the activity of the dorsal vessel of *R. prolixus*, both females and males, during feeding and evaluated the effects of meal temperature on the heart rate.

## MATERIALS AND METHODS

### 1. Insects

Experimental bugs came from our laboratory rearing stock at the Research Institute on Insects Biology at the University of Tours (France), where insects are kept under a 12:12 h light:dark regime and at 26±1°C, 60-70% relative humidity. Insects are fed weekly on heparinized sheep blood at 37°C using an artificial feeder (Núñez and Lazzari, 1990). Both 8 to 12. day-old unfed females and males of *Rhodnius prolixus* Stål (1859) (Heteroptera: Reduviidae: Triatominae) were used because of their relatively transparent dorsal cuticle beneath their wings, compared to larvae, which allowed a direct observation of the heart contractions without dissection. Wings were removed the day before the experiments, allowing the insect to heal and recover from a possible stress that could alter their heart rate. We considered a heartbeat as a contraction of the heart followed by a peristaltic wave starting at the abdominal end of the aorta (Martens and Chiang, 2010) (Movie 1). All experiments were performed in a room at 23°± 1°C and under dim-light to minimize stress.

### 2. Meal-temperature impact on the heart rate

#### 2.1 Heart rate at rest

By measuring the heart rate of insects at rest, we obtained a baseline for comparisons with heart rates recorded during feeding in both, females (n = 13) and males (n = 14). Each insect was fixed with a double-sided adhesive tape (∼ 2 × 2 mm) to a Petri dish in order to minimize movements during recordings. The dish was placed under a stereomicroscope fitted with a video-camera (Conrad, Hirschau, Germany) connected to a computer (Figure 1A). Heart activity was recorded using the Video Studio6 Ulead for further analysis of variations in the heart rate. For each experiment, five minutes of familiarization with the experimental setup were observed, followed by ten minutes of recordings for each insect.

**Figure 1.**
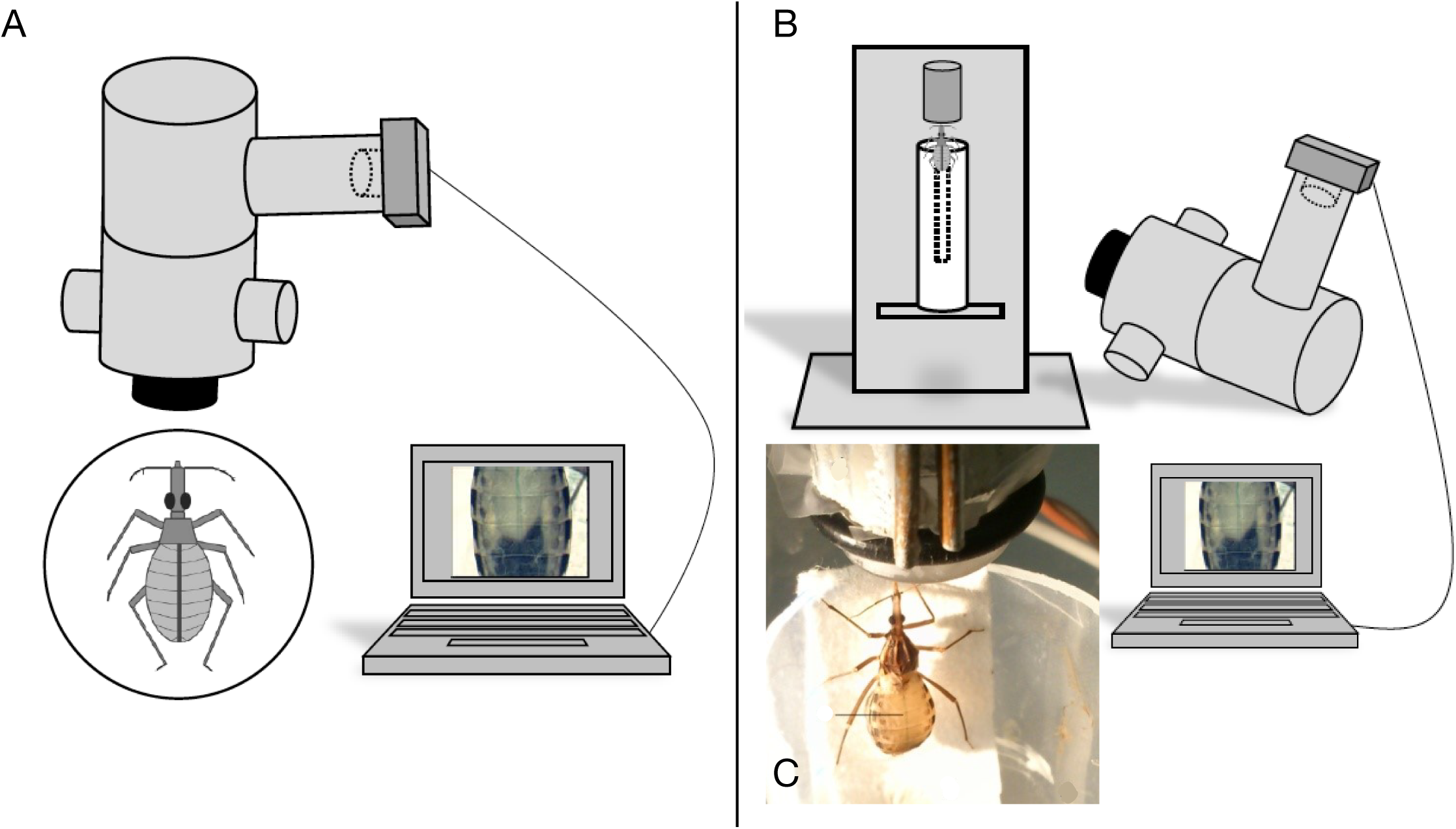
**A.** Experimental setup used for determining the heart rate at rest. The insect was fixed in a Petri dish. A camera mounted on a stereo microscope linked to a computer was used to record the heartbeats. **B.** Experimental setup used for heartbeat measurements during feeding on an artificial feeder. **C.** *Rhodnius prolixus* feeding on an artificial diet composed of saline solution which made the dorsal vessel (greenish line) visible through the insect cuticle.

#### 2.2. Heart rate during feeding

Insects were placed individually in a Falcon tube in which we cut a window to record the dorsal vessel contractions of the insect during feeding (Figure 1B). A strip of filter paper was placed in the tube, allowing the insect to reach the membrane of the artificial feeder placed above. In this position, the insect was not in contact with any warm surface, thus minimizing heat conduction (Figure 1C).

The meal consisted of *Rhodnius* saline solution (Maddrell, 1969). The transparency of this solution, compared to blood, improved the visibility of the dorsal vessel through the insect’s cuticle (Figure 1C). The saline solution was enriched with ATP (Sigma Aldrich, Saint-Louis, MO, USA) at 10-4M which is an efficient phagostimulant for *R. prolixus*, facilitating the meal intake (Smith and Friend, 1976).

The meal was provided to the insects with an artificial feeder equipped with a thermostat to maintain the meal at a specific temperature (*T*_*meal*_). Three meal temperatures were tested: 32°C (females n = 15; males n = 15), 37°C (females n = 16; males n = 18) and 42°C (females n = 19; males n = 19) (± 1 °C). *T*_*meal*_ = 32°C represents the minimal temperature at which *R. prolixus* considers the source as a potential meal and feeds naturally. *T*_*meal*_ = 37°C is in the range of the preferred meal temperature determined in this species (Wigglesworth and Gillett, 1934). *T*_*meal*_ = 42°C corresponds to a meal taken on birds whose body temperature are the highest among the hosts on which *R. prolixus* feeds on. These temperatures were previously successfully used to feed *R. prolixus* (Lahondère et al., 2017).

*Heart rate recordings.* Heart contractions were recorded during the feeding process. We considered that the meal began when the insect stopped moving, probing and as soon as we saw an abdominal expansion signaling that the gorging phase had already started. The end of feeding was determined when the insect withdrew its mouth-parts from the feeder. We compared heart rates of both, females and males, at rest with those of insects during feeding at different meal temperatures to evaluate the impact of the temperature on the heartbeat frequency.

### 3. Heart rate measurements and statistical analyses

For each group, we calculated the heart rate from five one-minute long sequences that were randomly selected along the 10 minutes recording time. Heartbeats were counted for each of these one-minute sequences. Heart rates (heartbeats/minute) were averaged within and across individuals. For measurements made during feeding, sequences were chosen with taking care of excluding the probing phase (Friend and Smith, 1971, 1977). After assessing the normality of the data distribution, a *two-way* ANOVA was performed using R (R Development Core Team, 2017) followed by a Tukey post-hoc test.

## RESULTS

First, we measured the heart rate at rest for both unfed females and males. Males exhibited a heart rate of 27.7± 4.1 b/min (mean ± S.E.M) and a mean of 28± 4.6 b/min was obtained for females (Figure 2). The statistical comparison of the heart rates of males and females did not show a significant difference (*t*-test, *p* = 0.83).

**Figure 2.**
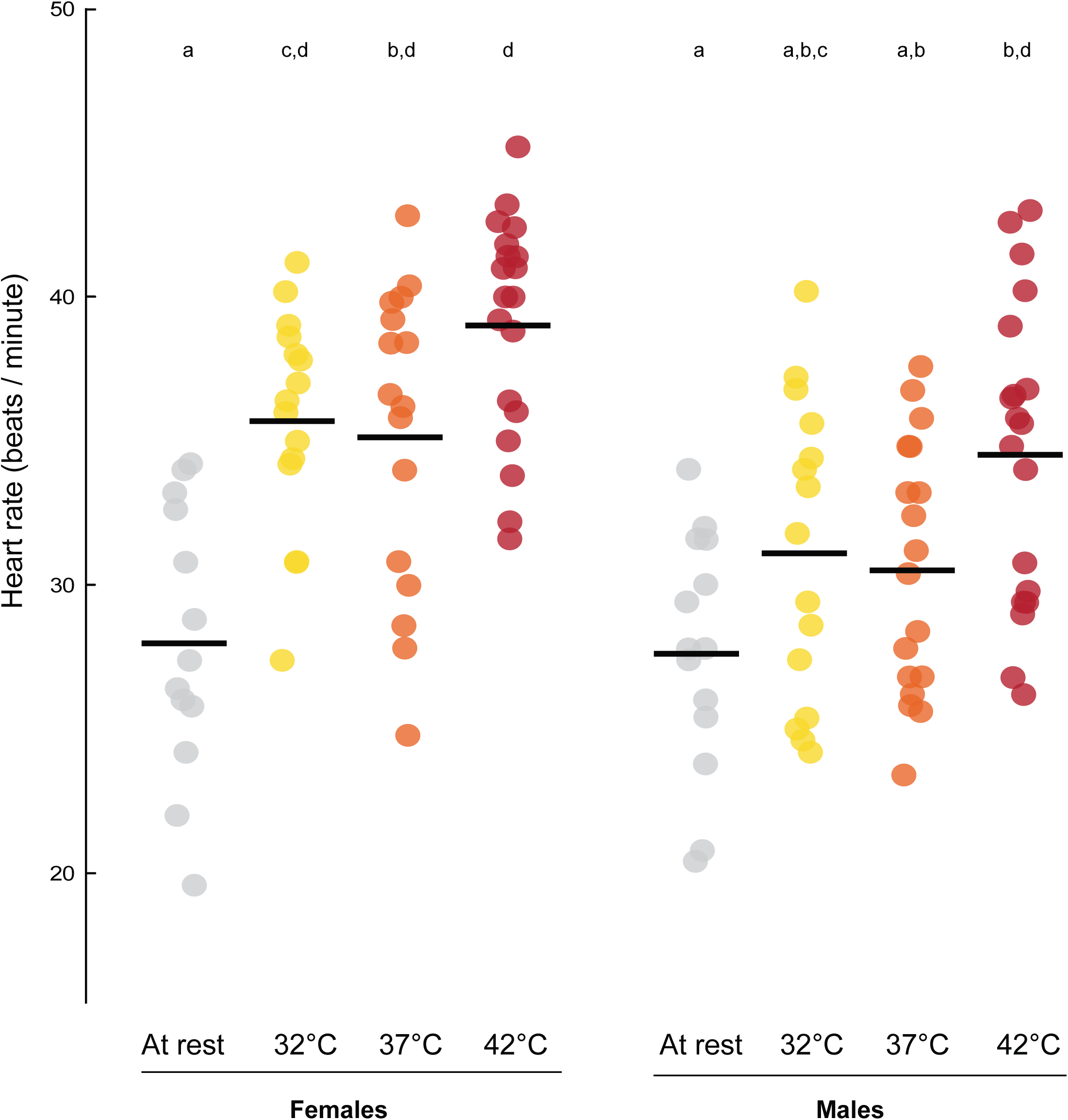
Heart rates of females and males *Rhodnius prolixus* bugs at rest (*T*_*a*_ = 23±1°C, in grey) and during feeding on meals at different temperatures (32°C in yellow, 37°C in orange, 42°C in red). Each set of dots represents an experimental group and each dot a single individual. Males groups: at rest (n= 14); *T*_*meal*_ = 32°C (n= 15); *T*_*meal*_ = 37°C (n= 18); *T*_*meal*_ = 42°C (n= 19). Females groups: at rest (n= 13); *T*_*meal*_ = 32°C (n= 15) *T*_*meal*_ = 37°C (n= 16) *T*_*meal*_ = 42°C (n= 19). Horizontal lines represent the mean. Different letters above boxes indicate statistical differences between groups (*P* < 0.05).

In a second time, we fed three groups of males and three groups of females, each at 32°C, 37°C or 42°C. A *two-way* ANOVA (factors: sex and temperature) revealed a significant effect of each of the two factors on the heart rate, but not a significant interaction between them (Table 1). We then performed a *Tukey post-hoc* test to further explore the effects of these factors on the insect heart rates. In both females and males, whatever the temperature of the meal, feeding led to an increase in the heart rate compared to rest (Figure 2). In males, the heart rate was only slightly higher at 32°C (31.2± 5.2 b/min) and 37°C (30.6± 4.4 b/min), but not significantly different to resting insects. Nevertheless, it was significantly higher when males were fed on saline solution at 42°C (34.6± 5.3 b/min, *p* = 0.001). In females, heart rates were significantly higher during feeding than at rest, for all three meal temperatures (35.8± 3.8 b/min at 32°C, 35.2± 5.3 b/min at 37°C and 39.1± 3.9 b/min at 42°C, *p* < 0.001 for all comparisons). Finally, we did not find any difference between females and males at rest or when fed at the three temperatures (Figure 2).

**Table 1.** *Two-way* ANOVA results for the analysis of the impact of the meal temperature and the sex on the heart rate of *Rhodnius prolixus*.

| Source of variation | DF | Sum of squares | Mean square | F-value | P-value |
| --- | --- | --- | --- | --- | --- |
| **Sex** | 1 | 458.3 | 458.3 | 21.403 | 9.41e-06 *** |
| **Temperature** | 3 | 1257.2 | 419.1 | 19.572 | 2.07e-10 *** |
| **Sex\*Temperature** | 3 | 94.0 | 31.3 | 1.463 | 0.228 |
| **Residual** | 121 | 2590.8 | 21.4 |  |  |
| **Total** | 128 |  |  |  |  |

## DISCUSSION

In *R. prolixus*, the active transport of cool hemolymph from the distal part of the abdomen to the head is essential for the counter-current heat exchanger to occur (Lahondère *et al.*, 2017). Even if our knowledge on hemolymph circulation in this species is limited, if cool abdominal hemolymph is not pumped by the heart and does not flow in the dorsal vessel at the adequate rate, we can expect that the countercurrent heat exchange occurring between the aorta and the oesophagus would not be efficient enough. Interestingly, it has been shown that *R. prolixus* bugs with a severed dorsal vessel do not present a heterothermic profile during blood-feeding, which underlines the important role of hemolymph circulation in thermoregulation during blood-feeding in this species (Lahondère *et al.*, 2017). During feeding, the anterior part of the head and the distal part of the abdomen are respectively the warmest and the coldest regions of the bug body and haemolymph circulation ensures this gradient to be kept. The experiments presented here provide evidence on the regulation of heart activity during feeding, in relation to the temperature of the meal. The heart rates measured in insects at rest (28± 4.6 b/min in females; 27.7± 4.1 b/min in males) are consistent with those obtained in the same species by Chiang *et al.* (33.3± 4.2 b/min) (Chiang *et al*., 1992) and Baehr and Baudry (1970) who found an average rate of 32 b/min. These studies also mentioned the variability in heart rates among individuals, from 28 to 41.1 b/min (Chiang *et al*., 1992) and from 20 to 40 b/min in the study of Baehr and Baudry (1970). This inter-individual variability was also noted in our study with heart rates ranging from 19.6 to 34.2 b/min in females and from 20.4 to 34 b/min in males. Feeding led to an increase of heartbeat frequency in *R. prolixus* in a temperature-dependent manner. We found that the mean heart rates of females were significantly higher at *T*_*meal*_ = 32°C and 37°C than at rest. Moreover, at *T*_*meal*_ = 42°C, the heart rates of both sexes were higher than at *T*_*meal*_ = 32°C and *T*_*meal*_ = 37°C. Interestingly, whatever the meal temperature, females exhibited higher heart rates than males. According to Jones (1964), the sex and the size of the individual make part of the factors that may influence the impact of temperature on the insect heart. In the case of *R. prolixus*, the fact that females are larger and take bigger meals (100.99± 22.18 mg) than males (63.29± 10.66 mg) could explain why they have a higher heart rate during feeding, given that the total amount of heat associated with the ingestion of the higher volume of fluid requires more efficient heat dissipation. We can thus hypothesize that insects increasing their heart rate would be able to eliminate the excess of heat at a higher rate. It is worth mentioning that in *Triatoma brasiliensis*, it has been shown that the volume of the cibarial pump is different between females and males and that females showed better feeding performance compared to males in regards of the ratio between the quantity of liquid ingested over the initial weight (Guarneri *et al.*, 2003).

In *R. prolixus*, very little is known about the hemolymph circulation compared to other insects (Hillyer, 2018) and much of the research effort has been devoted to its role for the circulation of factors involved in specific physiological processes, such as egg production (Chiang and Davey, 1990; Davey and Chiang, 1989), development and molting (Wigglesworth, 1934). Our work provides further evidence about the involvement of haemolymph circulation and the heart rate in coping with the thermal stress associated to feeding on the blood of warm-blooded vertebrates. It would be interesting to further explore how the temperature of the environment might affect *R. prolixus* heart rate and decouple the impact of the meal temperature and the subsequent effects of serotonin released during feeding for example.

In summary, the thermoregulatory ability of kissing-bugs depends on haemolymph circulation, as previously revealed by morphological studies and experiments in insects with a severed dorsal vessel (Lahondère *et al.*, 2017). The present study sheds additional light on the process, by unravelling the existence of a temperature-dependent modulation of heart activity associated with the temperature of the meal and underlines the interest on understanding the fine mechanisms behind heart rate regulation in kissing-bugs.

## Acknowledgments

We are indebted to Carole Labrousse for her technical support in *Rhodnius prolixus* rearing. We are very grateful to Clément Vinauger and Teresita Insausti for their valuable comments on the manuscript and helpful discussions. This work received financial support from the Agence Nationale de la Recherche (grant ANR-08-MIE-007, EcoEpi), the Centre National de la Recherche Scientifique, and the University of Tours (France).

## FIGURE LEGENDS

**Movie 1.** Video of a female *Rhodnius prolixus* focusing on its abdomen. Note the expansion of the dorsal vessel when hemolymph enters the heart and is then propelled along the aorta.

